# Phosphoproteomics reveals new insights into the role of PknG during the persistence of pathogenic mycobacteria in host macrophages

**DOI:** 10.1101/2021.01.19.427367

**Authors:** Seanantha S. Baros-Steyl, Kehilwe C. Nakedi, Tariq A. Ganief, Javan O. Okendo, David L. Tabb, Nelson C. Soares, Jonathan M. Blackburn

**Author notes:** **Correspondence**: Prof. Jonathan M Blackburn, Dr Nelson C Soares.

## Abstract

Pathogenic mycobacteria, such as *Mycobacterium tuberculosis*, modulate the host immune system to evade clearance and promote long-term persistence, resulting in disease progression or latent infection. Understanding the mechanisms pathogenic mycobacteria use to escape elimination by the host immune system is critical to better understanding the molecular mechanisms of mycobacterial infection. Protein kinase G (PknG) in pathogenic mycobacteria has been shown to play an important role in avoiding clearance by macrophages through blocking phagosome-lysosome fusion; however, the exact mechanism is not completely understood. Here, to further investigate the role of mycobacterial PknG during early events of macrophage infection, RAW 264.7 macrophage cell lines were infected with *M. bovis* BCG wild-type and PknG knock-out mutant strains. After proteolysis, phosphopeptides were enriched via TiO2 columns and subjected to LC-MS/MS to identify differentially phosphorylated peptides between the wild-type and PknG mutant infected macrophages. A total of 1401 phosphosites on 914 unique proteins were identified. Following phosphoproteome normalisation and differential expression analysis, a total of 149 phosphosites were differentially phosphorylated in the wild-type infected RAW 264.7 macrophages versus the PknG knock-out mutant. A subset of 95 phosphosites was differentially up-regulated in the presence of PknG. Functional analysis of our data revealed that PknG kinase activity reprograms normal macrophage function through interfering with host cytoskeletal organisation, spliceosomal machinery, translational initiation, and programmed cell death. Differentially phosphorylated proteins in this study serve as a foundation for further validation and PknG host substrate assignment.

**Importance:** Tuberculosis (TB) remains one of the leading causes of death from infection worldwide, due to the ability of *Mycobacterium tuberculosis* (*Mtb*) to survive and replicate within the host, establishing reservoirs of live bacteria that promote persistence and recurrence of disease. Understanding the mechanisms that *Mtb* uses to evade the host immune system is thus a major goal in the TB field. Protein kinase G is thought to play an important role in *Mtb* avoiding clearance by the host through disruption of macrophage function, but the underlying molecular mechanisms of this are not well understood. Here, our new phosphoproteomic data reveals that mycobacterial PknG substantially reprograms normal macrophage function through extensive PknG-mediated post-translational control of critical host cellular processes. These novel findings therefore considerably increase our knowledge of mycobacterial pathogenicity, including specific host cellular pathways that might be re-activatable through host-directed therapy, thereby restoring macrophage ability to eliminate *Mtb*.

## Introduction

Tuberculosis (TB) remains one of the leading causes of death from infection worldwide (1). The high prevalence of TB is due to the ability of *Mycobacterium tuberculosis (Mtb)* to survive and replicate within the alveolar macrophages, establishing reservoirs of live bacteria that promote persistence and recurrence of the disease. An estimated one-quarter of the global population harbours a latent TB infection (LTBI) (2), which, although asymptomatic, represent a reservoir for potential reactivation and transmission of TB (3).

Pathogenic mycobacteria suppress various host processes including phagosome-lysosome fusion, programmed cell death, and antigen presentation (4). In addition, *Mtb* promotes the activation of pathways involving mitogen-activated protein kinases (MAPKs), calcium signalling, and interferon-γ (IFN-γ) to weaken mycobactericidal responses (5, 6). Mycobacterial Ser/Thr protein kinases (STPKs) and phosphatases are known to play a role in interfering with host-signalling pathways during infection, thereby promoting intracellular survival (9–12).

The mycobacterial secretory protein kinase G (PknG) is a known virulence factor and contributes to the inhibition of phagosome-lysosomal fusion during early-stage infection (9). This is presumably a result of a kinase-mediated signalling mechanism through which PknG interacts/phosphorylates host protein substrates, thereby facilitating intracellular survival of the pathogen (9, 13). Experimental evidence shows that inactivation of PknG, either by gene disruption or chemical inhibition, results in lysosomal localisation and mycobacterial cell death in infected macrophages (9, 13). Although these processes are seemingly dependent on the kinase activity of PknG, the exact mechanism that PknG employs to establish a niche inside host macrophages remains unclear. Thus, understanding the mechanistic connection between PknG in such complex regulatory signalling networks and mycobacterial survival in macrophages requires a detailed view of the phosphorylation events taking place at a given time during infection.

Our previous *in vivo* study identified novel mycobacterial physiological substrates for PknG by comparing the phosphoproteome dynamics during growth in liquid culture of *M. bovis* BCG wild-type (WT) against the respective PknG knock-out (ΔPknG) mutant (14). The study revealed a new set of protein targets that were seen to be exclusively phosphorylated in *M. bovis* BCG WT and not in the ΔPknG mutant. Further validation of these initial results using PRMs and docking analyses allowed the identification of a set of mycobacterial proteins as novel physiological substrates for PknG (14).

Here, by employing a similar strategy, we aim to extend our knowledge of mycobacterial Ser/Thr/Tyr phosphorylation networks to *M. bovis* BCG-infected macrophages. Initial events during infection of host cells by pathogens can guide the overall course of infection and determine its eventual outcome. To understand host-pathogen interactions during this early stage of infection at the post-translational level, we analysed changes in the phosphoproteome of RAW 264.7 macrophage cells after infection with *M. bovis* BCG WT against those infected with the ΔPknG mutant. Through this work, we identify plausible protein phosphorylation mechanisms by which pathogenic mycobacterial PknG interferes with macrophage function to promote mycobacterial survival.

## Materials and Methods

### M. bovis BCG strains and bacterial culture

Bacterial strains used in this study were kindly donated by Professor Jean Pieters, University of Basel, Switzerland. The mutant (*M. bovis* BCG ΔPknG) was generated as described in Walburger et. al. (2004) (9). Mycobacteria were grown in 7H9 Middlebrook (BD, Maryland, USA) broth supplemented with 0.05% Tween 80, OADC (Becton Dickinson) and 0.2 % (v/v) glycerol. The cells were grown at 37 °C with continuous agitation (120 rpm). Growth was monitored daily by measuring the optical density (OD600) until the desired OD was reached.

### Culture conditions of RAW 264.7 macrophages

The murine macrophage cell line RAW 264.7 was used to study the host-pathogen interactions in this study. Murine macrophages have been used as a successful model to study host-pathogen interaction in *Mtb* studies. Our stocks were kindly donated by one of our collaborators from the University of Stellenbosch. Frozen stocks of the cell lines were defrosted at 37°C in a water bath. The cells in suspension were pelleted by centrifugation at 800 x g for 3 minutes. The cell pellets were washed twice with pre-warmed PBS and pelleted. The pellets were re-suspended in Dulbecco’s modified Eagle medium (DMEM) (Difco), supplemented with pyruvate and glutamine (Difco) and 10% (v/v) heat-inactivated foetal calf serum (FCS) (Sigma-Aldrich) (D10). The cells were then seeded into T25 tissue culture flasks (Lasec, South Africa) at 30% confluence and grown in a humidified (95%) incubator at 37°C with 5% CO_2_.

### Infection of macrophages with M. bovis BCG strains

Bacterial cells, at the desired OD based on the multiplicity of infection (MOI), were washed twice with PBS, and D10 was added to the cell pellet. Bacterial clumps were minimised using a water bath sonicator for 10 minutes. Bacterial cells were also passed through a needle 10 times using a 10 mL syringe, followed by gentle centrifugation (1000 x g for 5 min) to sediment clumps. Macrophage media was removed and washed twice with pre-warmed PBS and replaced with bacteria in D10 at MOI 1:4. Cells were incubated for 30 minutes uptake and washed five times with PBS. Fresh D10 was then added and incubated for a further 30 minutes and harvested. A modified RIPA buffer was added to the cells and agitated for 5 minutes on ice prior to treatment with endonuclease benzonase (Sigma).

### Tryptic digestion and phosphopeptide enrichment

In-solution digestion was carried out on 500 µg total protein. Briefly, proteins were denatured with 1 mM dithiothreitol (DTT) for 1 hour at room temperature with gentle agitation and alkylated with 5.5 mM iodoacetamide (IAA) for 1 hour in the dark. Proteins were pre-digested with Lys-C endopeptidase (Wako, Neuss, Germany) for 3 hours at 30°C before being diluted four times with HPLC-grade water to a final concentration of ~1.5 M urea. The diluted sample was then digested overnight with trypsin (1:100 ratio) at 30°C. The digestion was quenched with 1% trifluoroacetic acid (TFA) (Sigma Aldrich, St Louis, USA).

Digested peptides were desalted using in-house reverse-phase C18 chromatography (Millipore) in preparation for mass spectrometry (MS) analysis. Briefly, C18 columns were equilibrated twice with solvent B [80% acetonitrile (ACN) and 0.1% formic acid (FA)] added and centrifuged at 3000 x g for 30 seconds. Samples were washed twice with solvent A (2% ACN and 0.1%FA) before being added to the column. Peptides were eluted with solvent C (60% ACN and 0.1% FA) and dried down at room temperature in a SpeedVac vacuum concentrator (Savant). 50 µg of protein was reserved for proteome analysis and the rest enriched for phosphorylated peptides using a TiO_2_ phosphopeptide enrichment kit (ThermoFischer Scientific) according to the manufacturer’s instructions.

### Liquid chromatography with tandem mass spectrometry (LC-MS/MS analysis)

LC-MS/MS analysis was performed using the Dionex Ultimate 3500 RSLC Nano System (Thermo Fisher Scientific) coupled to a Q Exactive mass spectrometer (Thermo Fisher Scientific). After being desalted as described before, proteome and phosphoproteome peptides were resuspended in 15 µL of solvent A, of which 4 µL was loaded at 1 µg per injection into the LC-MS/MS system. Peptides were chromatographically separated using a 75 µm (ID), 25 cm column packed in-house with reversed-phase 3 µm Kinetex core-shell C18 resin (Phenomenex) at a flow rate of 400 nL/min at 40°C. The gradient consisted of 2% solvent B (0.1% FA, ACN) increased to 8% solvent B over 2 minutes, followed by increasing to 23% solvent B over 80 minutes, followed by a washout at 80% solvent B for 10 minutes.

MS1 spectra were acquired between 300-1750 Thompson at a resolution of 75000 with an AGC target of 3 × 106 within 250 ms. Using a dynamic exclusion window of 90 sec, the top 10 most intense ions were selected for higher-energy collisional dissociation (HCD) fragmentation with an NCE of 28. MS2 spectra were acquired at a resolution of 17500 and a minimum AGC of 1 × 103 within 80 ms.

### Protein and phosphopeptide identification

Raw files were processed in MaxQuant version 1.6.14.0. MS/MS spectra were searched against the *Mus musculus* reference proteome downloaded 7 July 2020 from UniProt (http://www.uniprot.org/proteomes/UP000000589) containing contaminants. Maxquant’s built-in Andromeda search algorithm (15) was used to map spectra to the reference proteome with mass tolerance for precursor and fragment ions set at 4.5 ppm and 20 ppm, respectively. The match-between-runs functionality was enabled, by which peptide identifications can be transferred across raw files based on accurate retention time and mass-to-charge ratio (m/z). The identification false-discovery rate (FDR) was estimated using a target-decoy database. Carbamidomethylation of cysteine residues was specified as a fixed modification for all groups. Variable modifications considered were oxidation of methionine and protein N-terminal acetylation for the proteome files, with the addition of phosphorylation of serine, threonine, and tyrosine (Ser/Thr/Tyr) residues for the phosphoproteome files. Trypsin and Lys-C were selected as digestion enzymes and two missed cleavages were allowed.

### Statistical processing & bioinformatics analysis

The Proteus package (16) in R was used for evidence data exploration, quality checks, and visualisation at the peptide level. Perseus version 1.6.14.0 software (17) was used for quality control, statistical processing, and protein annotation. Protein identifications were filtered for those only identified by site, reverse database hits, and potential contaminants. The log2-transformed protein abundance values were then filtered to contain a minimum of three valid values among the four biological replicates per condition. A two-sample t-test with a Benjamini-Hochberg (BH) FDR threshold of 0.05 was performed to detect differentially abundant proteins.

Phosphopeptide identifications were filtered for reverse hits, potential contaminants, and a localisation probability >0.75. The intensity values were log2-transformed before filtering for valid values. The phosphoproteomic intensities were normalised to the proteome intensities by subtraction to account for protein abundance differences and then filtered to contain a minimum of three valid values among the four biological replicates per condition. A median normalisation was also performed across the data set, followed by a two-sample t-test with a BH FDR threshold of 0.05. The identified differentially phosphorylated peptides were functionally annotated and filtered for unique entries.

### Functional enrichment and network analysis

To identify pathways altered by PknG during mycobacterial infection, we assessed the complement of differentially abundant proteins and phosphoproteins for enriched Kyoto Encyclopaedia of Genes and Genomes (KEGG) pathways using the enrichment analysis tool of the stringApp plug-in within Cytoscape (18, 19) with FDR <0.01. The *Mus musculus* genome was used as the background. Functional enrichment is based on over-representation analysis using hypergeometric tests. Similarly, the complement of differentially phosphorylated proteins and the subset of differentially up-regulated phosphoproteins were used to retrieve enriched Gene Ontology (GO) terms using the stringApp. The enrichment results were filtered for redundancy with a cut-off of 0.25 and GO terms and KEGG pathways identified with less than three proteins were ignored.

Visualisation of the functional enrichment networks was performed using the clusterProfiler package (20). Briefly, the genome-wide annotation library (org.Mm.eg.db) for *Mus musculus* was downloaded from Bioconductor (www.bioconductor.org). UniProt IDs of the differentially phosphorylated proteins were converted to Entrez Gene IDs using org.Mm.eg.db as the organism database. We achieved 90.28% mapping rate of our proteins to the annotation database. Gene-concept networks were generated using the GO enrichment analysis results.

## Results

### Intracellular survival of mycobacteria in RAW 264.7 macrophages

We monitored the survival dynamics of*M. bovis* BCG WT compared to the ΔPknG mutant, judged by measuring colony-forming units (CFUs) at five-hour intervals post-infection (Figure 1). Our results are in agreement with the work of Walhburger et al. (2004) (9) on which our present study is based. The BCG WT strain displayed a two-fold increase over 24 hours post-infection whilst the null mutant’s growth was controlled by the macrophages.

**Figure 1.**
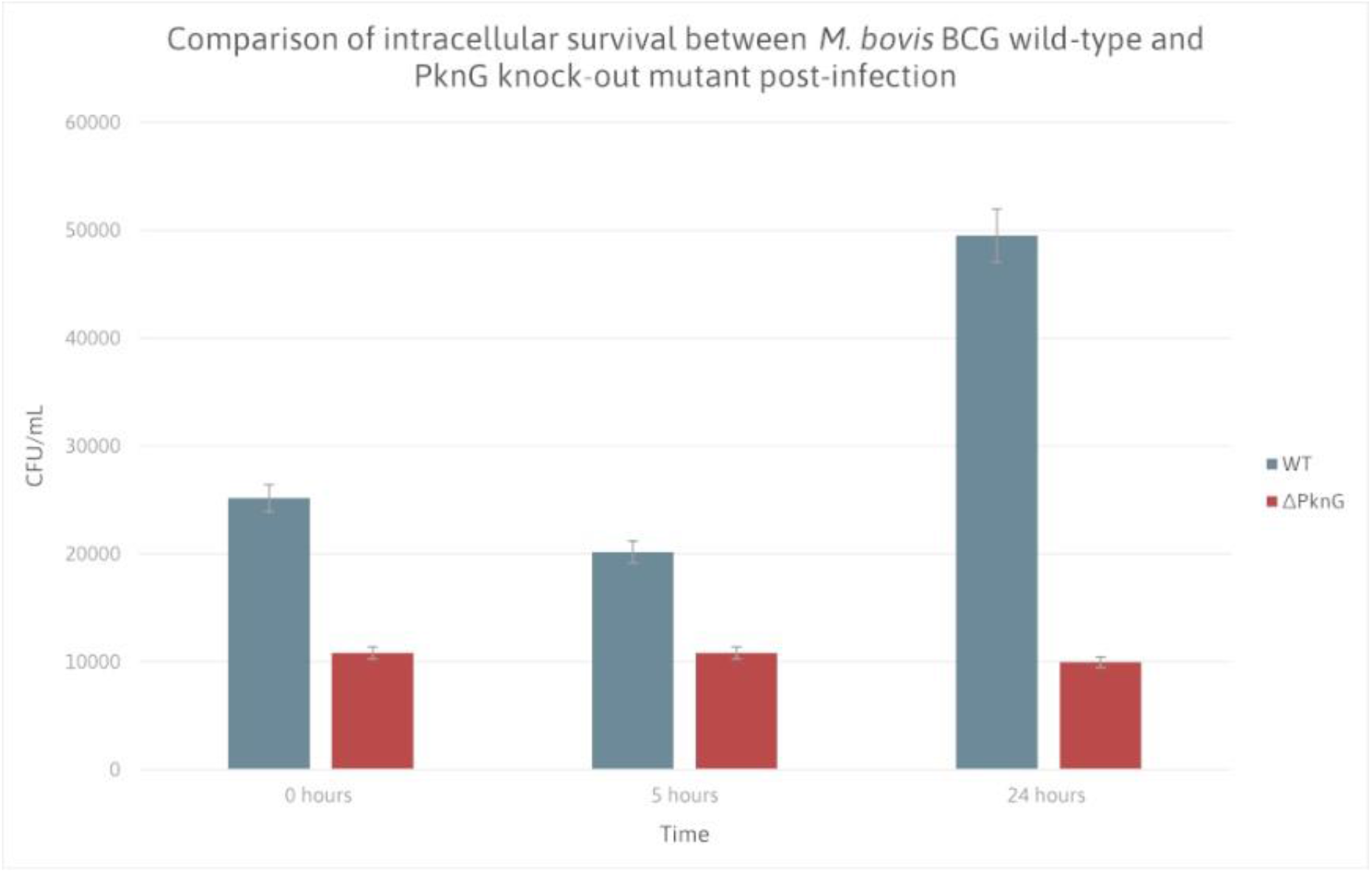
Survival of *M. bovis* BCG WT and ΔPknG mutant in macrophages at different time points post-infection. Histogram showing the bacterial growth of the different mycobacterial strains in RAW 264.7 macrophages at 0 hours, 5 hours, and 24 hours post-infection.

### Proteomic analysis of infected RAW 264.7 macrophages with mycobacterial strains

Proteomic experiments were run in parallel to the phosphoproteomic workflow (Figure 2). The evidence data was used to assess the quality of peptide identifications (Figure S1). After initial filtering, the proteomic experiment identified a total of 3441 protein groups. The log2-transformed label-free quantification (LFQ) protein abundance values were then filtered to contain a minimum of three valid values in each group, resulting in 1317 protein groups remaining. After differential analysis, 619 proteins were considered to have significantly different abundance levels between the *M. bovis* WT and ΔPknG samples (Table S1). Of these, 497 host proteins were up-regulated (Table S2).

**Figure 2.**
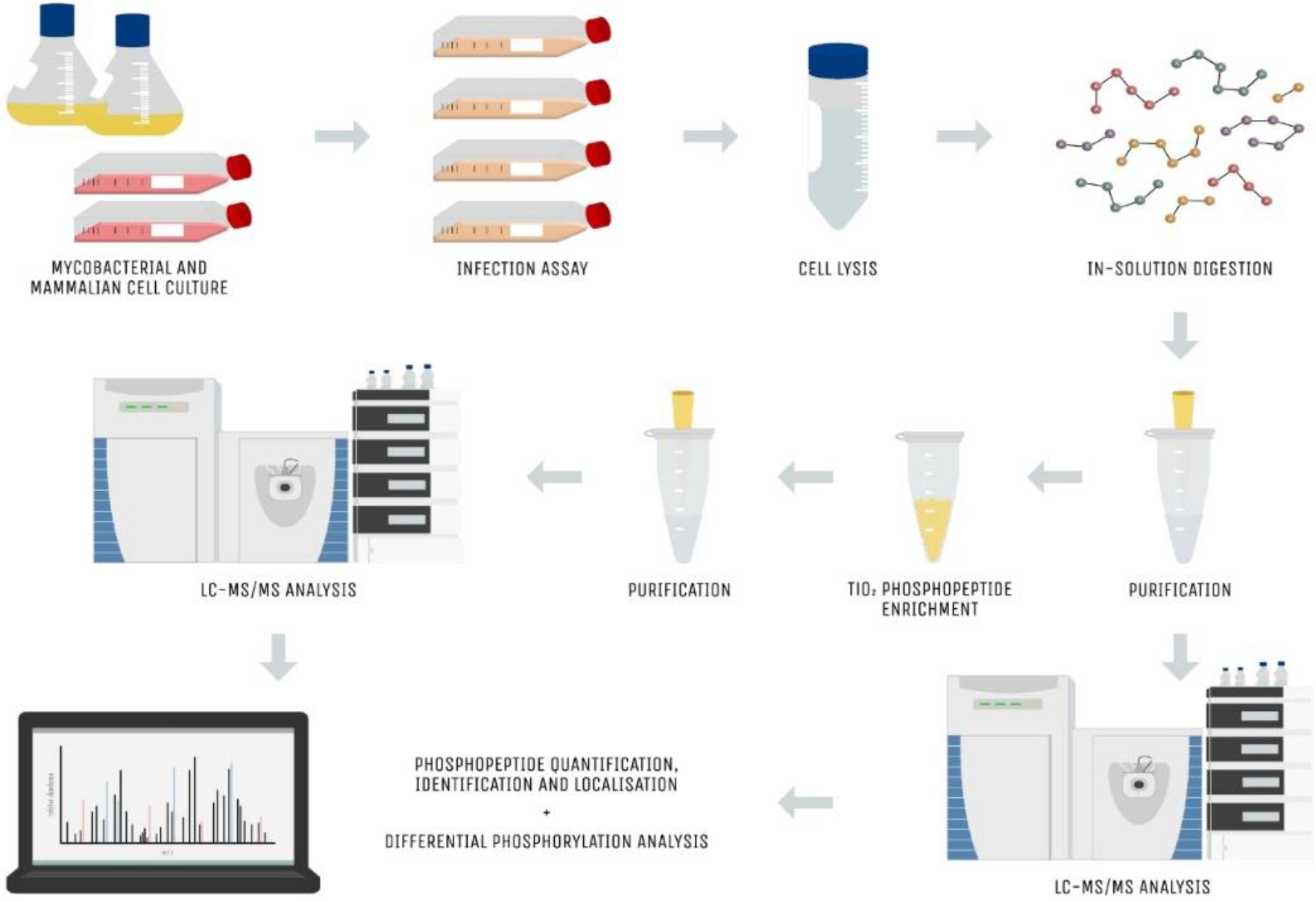
Proteomic and phosphoproteomic workflow used in this study. *M. bovis* BCG wild-type and ΔPknG strains were grown until mid-log phase and RAW 264.7 cells were cultured in D10 media until confluent. Infection assays were carried out at MOI 1:4 with a one-hour incubation. The RAW 264.7 cells were washed and lysed in the presence of protease and phosphatase inhibitors, followed by in-solution digestion with Lys-C and Trypsin. The samples were purified using reverse-phase C18 chromatography and an aliquot reserved for proteome analysis. Phosphopeptides were enriched using TiO_2_ phosphopeptide enrichment kit, followed by another clean-up step. LC-MS/MS data acquisition was then performed on an HPLC coupled to a Q Exactive mass spectrometer. The raw data files were processed using MaxQuant and analysed using Perseus and other software packages.

### Identification of PknG-mediated quantitative phosphoproteomic changes in the RAW 264.7 macrophages during infection

To identify potential host targets of mycobacterial PknG, we employed a label-free quantitative phosphoproteomics approach to study global phosphorylation dynamics associated with PknG during *M. bovis* BCG macrophage infection. Four biological replicates per experiment were analysed to increase the phosphoproteome coverage. We considered proteins phosphorylated in at least three of the four replicates per condition that displayed a statistically significant increase in phosphorylation in the WT as candidate host proteins phosphorylated via the direct or indirect action of PknG.

A total of 1401 phosphosites (p-sites) were identified. After removing known contaminants and filtering low-quality data points as previously described, 1371 highly confident phosphopeptides mapped to 914 phosphoproteins were taken for further analysis. The distribution of phosphoserine (pS), phosphothreonine (pT) and phosphotyrosine (pY) sites was 85.3%, 13.4%, and 1.3%, respectively. Moreover, the majority of the peptides were singly phosphorylated (95.1%), whereas approximately 4.9% were multiply phosphorylated. The mass spectrometry proteomics and phosphoproteomics data have been deposited to the ProteomeXchange Consortium (http://proteomecentral.proteomexchange.org) via the PRIDE partner repository with the dataset identifier PXD019365 (21, 22).

The quantification of phosphorylated proteins is important to our understanding of the resulting biological effects. To ensure that these differentially regulated phosphopeptides were not because of changes at the proteome level, the phosphoproteomic intensities were normalised to the proteome intensities. The normalised log2-transformed intensities of the RAW 264.7 cells infected with *M. bovis* BCG WT and ΔPknG mutant were then compared using a two-sample t-test with a BH FDR <0.05. A total of 149 peptides were significantly differentially phosphorylated (Table S3). Of these, the 95 p-sites were differentially up-regulated in the presence of mycobacterial PknG (Table S4) with 44 phosphoproteins differentially up-regulated with log2 fold-change (log2FC) >1 (Table 1). A volcano plot (Figure 3B) revealed significantly differentially phosphorylated peptides with FDR <0.05. Hierarchical clustering of the differentially up-regulated phosphopeptides with log2FC >1 (Figure 3C) was performed to assess the biological variability between samples. Figure S2 illustrates the relative fractions of pS, pT and pY observed among the differential and non-differential p-site, peptide, and protein identifications.

**Figure 3.**
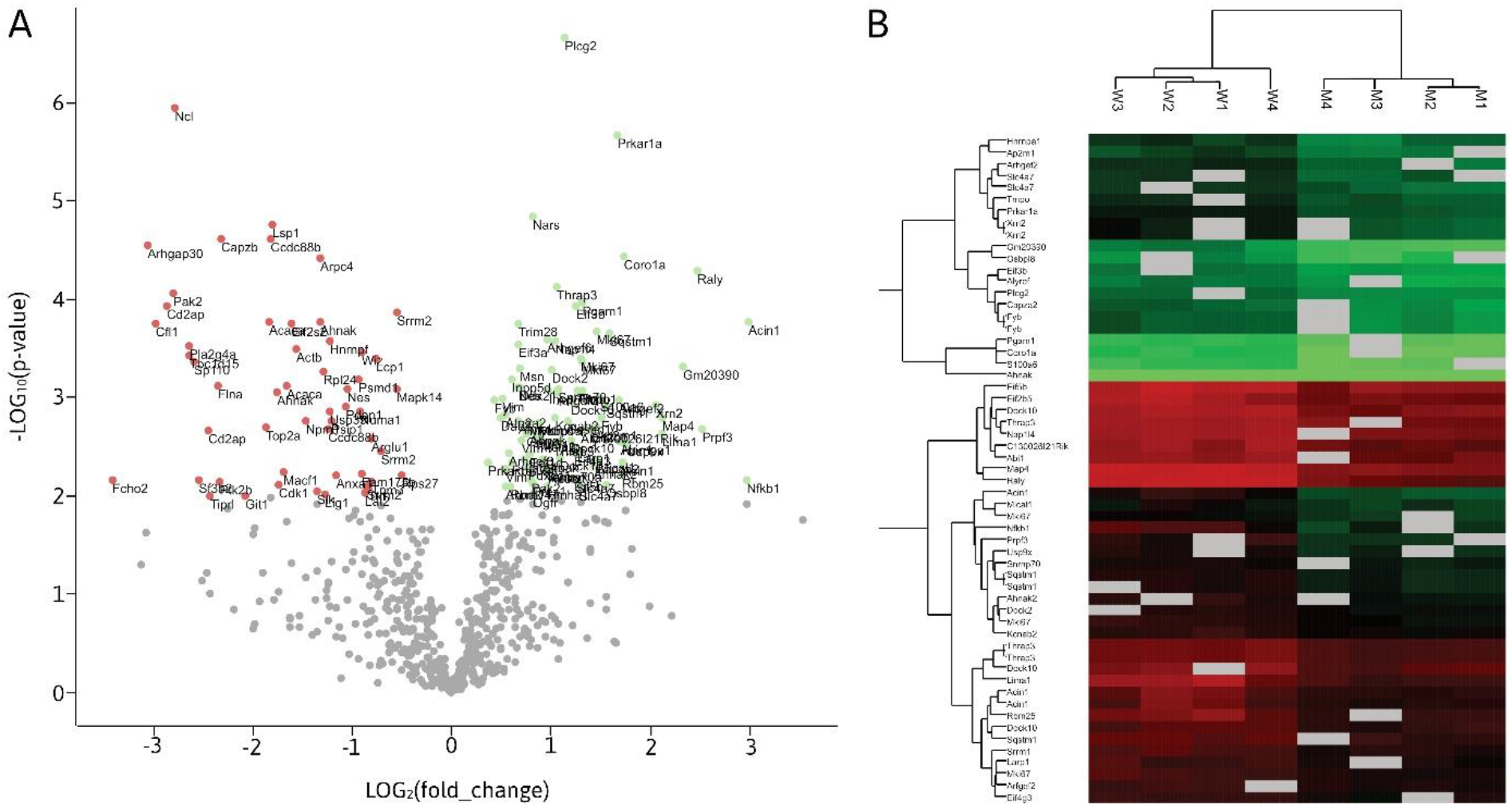
Analysis of quantified phosphopeptides. (A) Volcano plot highlighting the significantly differentially phosphorylated peptides. Scatter plot showing the phosphopeptide log2 fold-change (WT/ΔPknG) plotted against the -log10(p-value) highlighting the differentially phosphorylated peptides (two-sample t-test with BH FDR <0.05). Down-regulated and up-regulated phosphopeptides in the presence of PknG are indicated in red and green, respectively. **(B) Hierarchical clustering analysis of differentially up-regulated phosphopeptides in the presence of PknG.** Heatmap showing the abundance levels and clustering of the significantly differentially unregulated phosphopeptides (two-sample t-test with BH FDR <0.05; log2FC ≥1 WT/ΔPknG). Phosphopeptide intensities up-regulated are coloured green while down-regulated are coloured red. Row headings represent gene names and column headings represent samples.

**Table 1.**
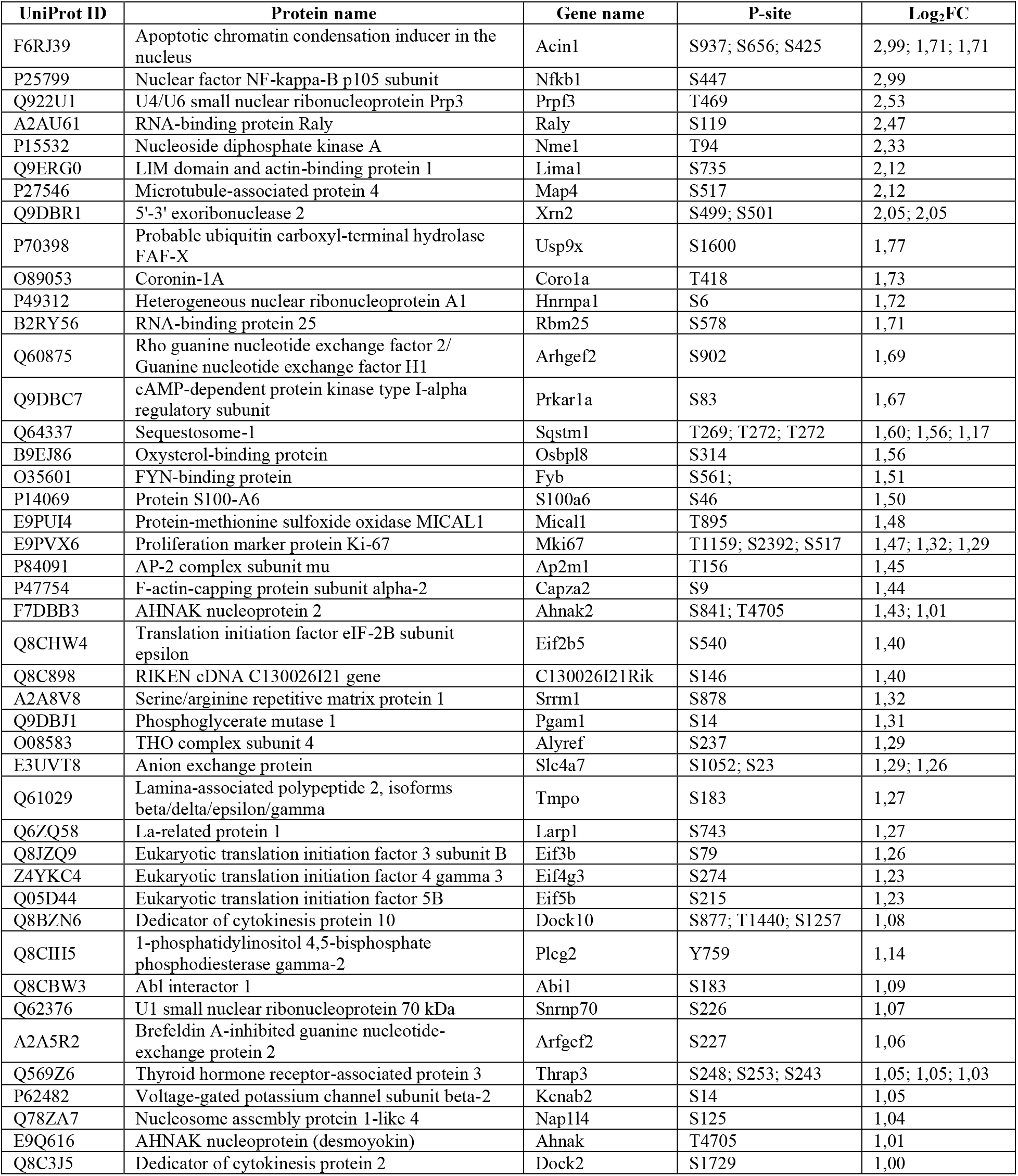
List of differentially regulated host phosphoproteins with a minimum two-fold increase in phosphorylation in *M. bovis* BCG WT respective to the ΔPknG mutant during infection.

### Functional annotation of host proteins differentially abundant proteins and phosphoproteins in the presence of PknG

Functional enrichments in our data were retrieved for GO terms and KEGG pathways and filtered for redundancy. A functional enrichment network was built for the significantly differentially abundant proteins and phosphoproteins using the clusterProfiler package (Figure 4). Enriched KEGG pathways of the differentially abundant proteins (Figure 5A) reaffirm PknG’s role in the regulation of host metabolism (23) and the phagosome-lysosome interaction (9, 24, 25) to control mycobacterial virulence. These results also propose a role for PknG in the manipulation of host translational, proteasomal, and spliceosomal machinery to promote intracellular survival of the bacilli. Among the functionally enriched KEGG pathways of the differentially phosphorylated proteins (Figure 5B), Fc gamma R-mediated phagocytosis, regulation of actin cytoskeleton, and endocytosis are consistent with previous literature surrounding the intracellular role of mycobacterial PknG during infection (9, 13, 25). However, observing enriched pathways such as RNA transport, the spliceosome, and DNA replication in our dataset suggests that PknG influences a wide range of host cellular processes.

**Figure 4.**
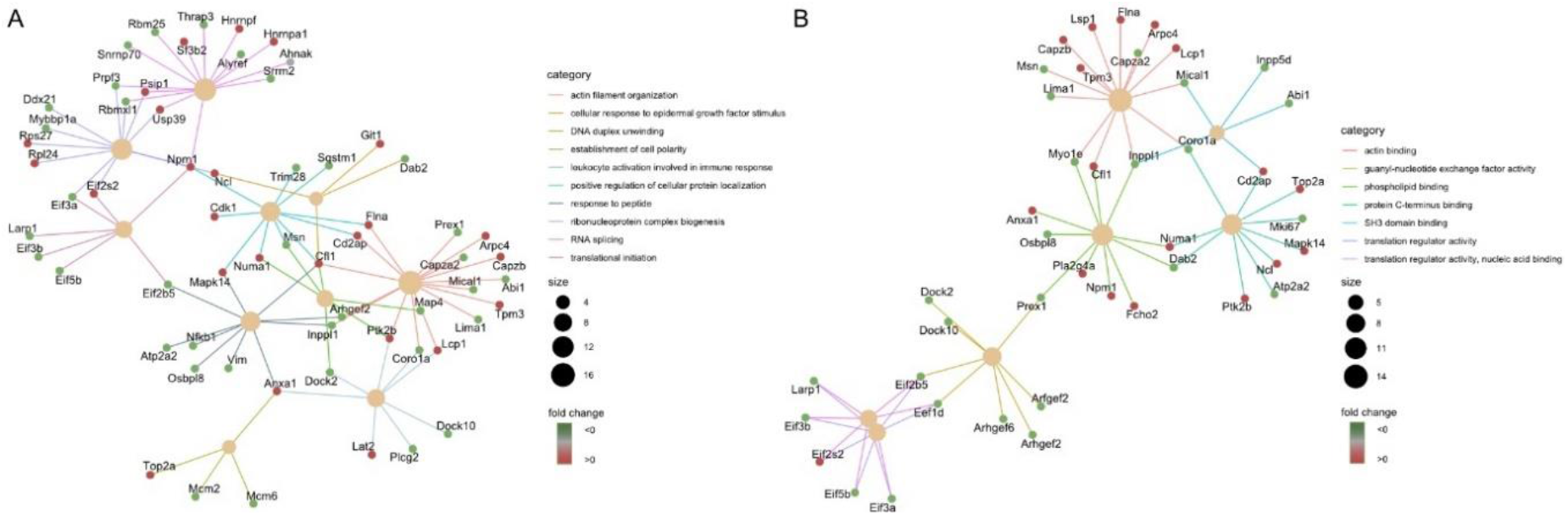
Functional enrichment network of differentially phosphorylated proteins in the presence of mycobacterial PknG associated with up to 10 most significant GO (A) biological processes and (B) molecular functions. Nodes are coloured according to fold change and sized according to the number of associated proteins.

**Figure 5.**
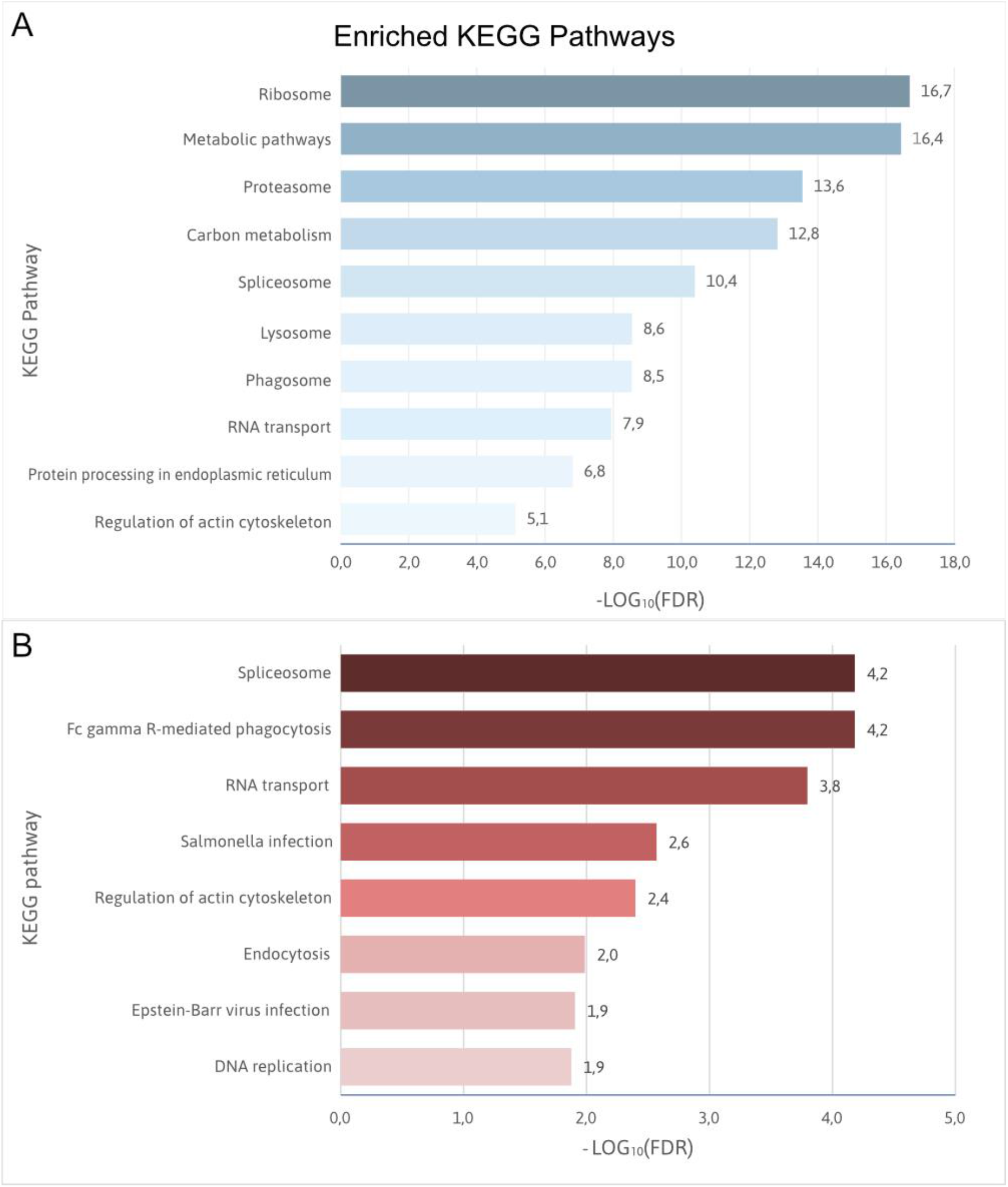
Significantly enriched KEGG pathways of differentially abundant (A) proteins and (B) phosphoproteins. Bar charts ranking the significant Kyoto Encyclopaedia of Genes and Genomes (KEGG) pathways retrieved after functional enrichment analysis of the differentially regulated proteins and phosphoproteins in the presence of PknG during mycobacterial infection.

Cellular component GO enrichment of the differentially up-regulated phosphoproteins (Figure 6A) shows that the majority of their localisation is in plasma membrane-bounded cell projections, the spliceosomal complex, cytoplasmic region, and actin cytoskeleton. Complementary to this, Figure 6B shows that the highly enriched biological processes are mRNA processing, translational initiation, regulation of cellular processes, and actin cytoskeleton organisation. Binding was observed as the most enriched molecular function among the differentially phosphorylated proteins (Figure 6C) and describes molecule interactions which are dominated by cytoskeletal protein binding and nucleic acid binding. Moreover, we see the enrichment of guanyl-nucleotide exchange factor (GEF) activity, which refers to the exchange of guanyl-nucleotides associated with a guanosine triphosphatase (GTPase).

**Figure 6.**
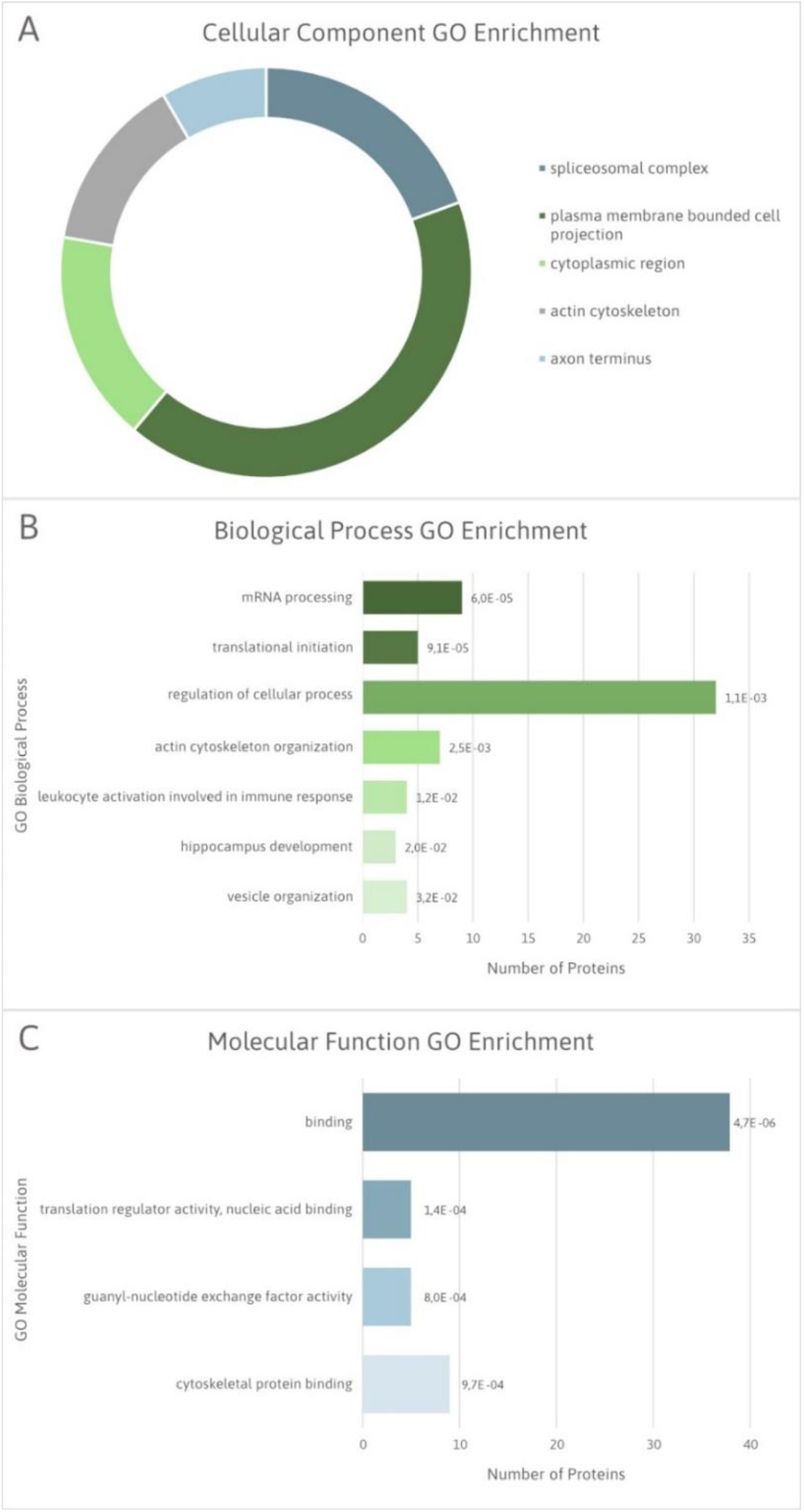
Significantly enriched GO terms of candidate host macrophage substrates of mycobacterial PknG. **(A)** Pie chart illustrating the significantly enriched cellular component GO terms of the candidate host PknG substrates. **(B)** Bar chart ranking of the significantly enriched biological process GO terms, and **(C)** molecular function GO terms of the candidate host PknG substrates. Data labels represent FDR values for each enriched GO term.

Notably, the Rho family GTPases regulate the integrity of the actin cytoskeleton (26). GTPases, RhoA, Rac1, and Cdc42, participate in the regulation of crucial processes that are dependent on the actin cytoskeleton such as cytokinesis, transcriptional activation, phagocytosis, morphology, and migration (27–30). The activation state of Rho GTPases is governed by the balance between the activities of GEFs and GTPase-activating proteins (GAPs) (31). Unsurprisingly, four GAPs and six GEFs were differentially phosphorylated in our data and serve to validate our findings.

Following from this, functional enrichment networks of the differentially up-regulated phosphoproteins highlighted subsets of proteins that appear to be interlinked and associated with actin filament organisation, negative regulation of cytoskeleton organisation, response to peptide, regulation of cellular amide metabolism and translational initiation (Figure 7A). Expectedly, the molecular functions that emerge as interlinked include actin binding and SH3 binding, as well as GEF activity and translation regulatory activity (Figure 7B). This highlights mycobacterial PknG’s ability to manipulate several key cellular functions during infection of host macrophages.

**Figure 7.**
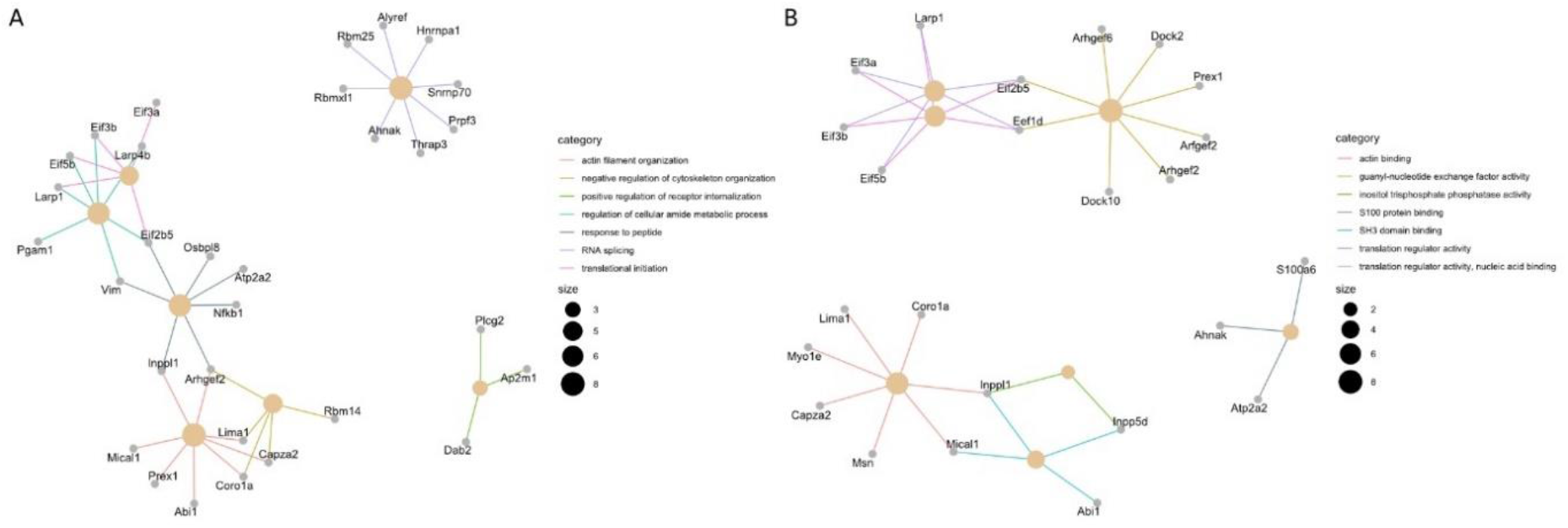
Functional enrichment network of differentially up-regulated phosphoproteins in the presence of PknG associated with significant GO (A) biological processes and (B) molecular functions. Nodes are sized according to the number of associated proteins.

## Discussion

PknG is thought to play an important role in reprogramming macrophage function to aid the survival of tubercle bacilli. Studies have demonstrated that macrophages harbouring *M. bovis* BCG strains failed to induce phagosome maturation, while the mutant lacking PknG was successfully eliminated (9, 32). This prompted us to decipher the host mechanisms exploited by PknG to promote the pathogenesis of TB, by identifying the host proteins phosphorylated by PknG after *M. bovis* BCG has been phagocytosed. To this end, we carried out a quantitative label-free phosphoproteomic approach that we, and others, have shown to be a high-throughput platform to identify targets of protein kinases (33–36). The functions of the differentially phosphorylated proteins identified in this study shed new light on the mechanisms that PknG uses to regulate and reprogram host-signalling pathways that enable the bacterium’s survival, as discussed below.

### PknG mediates the regulation of cytoskeletal organisation

The cytoskeleton is a three-dimensional network of proteins, including actin filaments, microtubules (MTs), and intermediate filaments (IFs) (37, 38). Actin cytoskeletal rearrangement promotes numerous events that are beneficial to intracellular pathogens, including internalisation of bacteria, altered vesicular trafficking, actin-dependent motility, and pathogen dissemination (39). A few groups have explored the role of the actin cytoskeleton during *Mycobacterium* late phases of phagocytosis. Guérin & De Chastellier (2000) showed that *Mycobacterium avium* disrupts the macrophage actin filament network, highlighting the target for the bacterium that allows sustained intracellular survival (40). Anes et al. (2003) demonstrated that in contrast to non-pathogenic mycobacteria, pathogenic *Mtb* prevents actin polymerisation on phagosomal membranes (41). We have identified six phosphoproteins that we believe are indicative of disruptive cytoskeletal remodelling in the presence of PknG.

### Rho GTPase-mediated cytoskeletal remodelling

As previously mentioned, GEFs and GAPs modulate Rho GTPase activity and ultimately coordinate cytoskeleton remodelling (26, 31). GEF-H1 is a unique GEF that associates with microtubules and is known to mediate the crosstalk between MTs and the actin cytoskeleton, which may contribute to the phagocytosis of bacteria by macrophages (42). The activity of GEF-H1 is suppressed when it is bound to the dynein motor complex on MTs (43, 44) and activated when released from MTs upon the intracellular binding of bacterial effectors (45, 46). A recent study showed that the expression of GEF-H1 was increased via the MAPK signalling pathway during mycobacterial infection and silencing of GEF-H1 inhibited macrophage-mediated mycobacterial phagocytosis and elimination c

Various studies have shown that GEF-H1 is both positively and negatively regulated via phosphorylation. For instance, it has been reported that GEF-H1 phosphorylation in the C-terminal portion on S885 and S959 contributes to holding GEF-H1 in a catalytically inactive configuration, thus inhibiting its GEF activity (47, 48). In contrast, phosphorylation that occurs between the PH and coiled-coil domain of GEF-H1 at S151 and T678 causes dissociation from microtubules (49, 50). This is followed by stimulation of RhoA activation, resulting in the formation of focal adhesions and stress fibres, thereby enabling actin cytoskeleton remodelling.

We identified phosphorylation of GEF-H1 at S902, which falls within the C-terminal region and should correspond with negative regulation of GEF activity. This suggests that phosphorylation of GEF-H1 in the presence of PknG hinders RhoA activation, thereby contributing to the disruption/uncoupling of actin stress fibres and focal adhesions. Such modification to the organisation and structure of the host cytoskeleton would promote the survival and growth of the bacterium in host macrophages, consistent with the GEF-H1 silencing studies discussed above, consistent with the GEF-H1 silencing studies discussed above (46).

The Rac1/Cdc42 GEF, ARHGEF6, plays certain roles in some immune cells, including neutrophils and T cells. Importantly, ARHGEF6 is required for the activation of Ser/Thr-protein kinase PAK 2 (PAK2) and subsequent LIM-kinase (LIMK)-dependent inactivation of cofilin. Cofilin-1 (Cfl1) was significantly dephosphorylated in the presence of PknG in our dataset. ARHGEF6 was found to be differentially phosphorylated at S663 and S703 in this study and the phosphorylation of PAK2 at S2 and S55 were significantly up-regulated and down-regulated, respectively. Collectively, the ARHGEF6/Rac1/PAK2/LIMK/cofilin signalling module limits actin turnover by confining both lamellipodial actin polymerisation and depolymerisation, enables polarized, stable lamellipodia formation, and promotes focal complex assembly in lymphocytes (51). Since phosphorylation of Cfl1 suppresses its activity, our study indicates enhanced actin depolymerisation and disruption of actin stress fibres and focal adhesion complexes in the presence of PknG. This also suggests that phosphorylation of ARHGEF6 at S663/703 has an inhibitory effect on its Rac1 GEF activity.

Similarly, DOCK2 activates Rac-dependent actin polymerization and contributes to T and B cell migration (52, 53). Dendritic cells defective in DOCK2 exhibit impaired endocytosis of soluble antigens and phagocytosis of insoluble antigens and larger particles (54). Moreover, DOCK10, a Rac1/Cdc42 GEF, is known to affect cell morphology, spreading, and actin cytoskeleton protrusions of adherent HeLa cells (55). Together, phosphorylation of these GEFs reaffirms the role that PknG plays in disrupting cytoskeletal organisation during infection.

Furthermore, LIM domain and actin-binding protein 1 (Lima1, also known as EPLIN) is involved in actin cytoskeleton regulation and dynamics. Lima1 cross-links and bundles actin filaments, thereby stabilising actin stress fibres. Lima1 inhibits Arp2/3 complex-mediated branching nucleation of actin filaments, thus controlling actin filament dynamics by stabilising actin filament networks (56). Phosphorylation of the C-terminal region of Lima1 inhibits its actin-binding activity (57). Thus, the increased phosphorylation of Lima1 in the presence of PknG contributes to the reduced stability of actin stress fibres and increased actin filament depolymerisation, as observed with GEF-H1.

### PknG inhibits phagosome maturation

Mammalian nucleoside diphosphate kinase (NDPK) proteins participate in the regulation of multiple cellular processes via enzymatic and non-enzymatic functions. Their primary enzymatic function is to catalyse the conversion of nucleoside diphosphates (NDPs) into nucleoside triphosphates (NTPs) (58). Here, NDPK-A [also known as NME1, NM23-H1/NM23-M1 (human/mouse homolog)] was identified with differential phosphorylation on T94 in the presence of PknG. Mounting evidence suggests that NDPK-A interacts with and affects various components and regulators of the cytoskeleton, including actin-binding proteins, IFs, and cytoskeletal attachment structures (59–66). Garzia et al. (2008) demonstrated that increased phosphorylation of NDPK-A promotes cell motility in breast cancer cells (67). Importantly, NDPK-A possesses GAP activity and negatively regulates Rho-Rac signalling through the inhibition of Rac1 activity (68, 69). NDK-A silencing increased Rac1 signalling and MAPK/SAPK (stress-activated protein kinases) activation (60).

A recent study showed that NDPK-A is a phagocytosis promoting factor and acts in a complex with Dynamin (DYN-1) on phagosomal surfaces, during engulfment and the early stages of phagosome maturation (66). NDPK-A depletion in macrophages led to decreased phagocytic capacity (66). Furthermore, NDPKs are known to be secreted by mycobacteria, including *Mtb* and *M. bovis* BCG (70, 71). *In vitro* analyses demonstrated that mycobacterial NDPK possesses GAP activity towards Rho GTPases, Rab5 and Rab7, which regulate endosomal trafficking in macrophages. It is suggested that mycobacterial NDPK disrupts the maturation and lysosome fusion of phagosome-containing pathogenic mycobacteria by inactivating Rab5 and Rab7 (72, 73).

Similarly, when bacteria are phagocytosed by antigen-presenting cells, the phagosome acquires early endosomal protein markers, such as early endosomal antigen 1 (EEA1) and Rab5, which are gradually replaced with Rab7 during the maturation of the phagosome (74, 75). It has been reported that phagosomes containing live pathogenic mycobacteria do not acquire Rab5 due to the transient recruitment and active retention of coronin-1A (also known as tryptophan aspartate coat protein (TACO) or p57) (76). Hence, stable phagosomal association with coronin-1A results in the inhibition of phagosomal maturation (77).

We identified the host F-actin-binding protein coronin-1A to be differentially phosphorylated on T418. Upregulation of T418 was also previously observed during the first 10 minutes after lipopolysaccharide (LPS) and Pam3Cys stimulation of macrophages (78). Notably, phosphorylation of coronin-1A deregulates its association with F-actin, which, in turn, facilitates early phagosome formation (50). Schuller et al. (2001) also found that *Mtb* inhibits lysosome formation by increasing the expression of coronin1-A on the phagosomal membrane (79). Together, our results suggest that PknG phosphorylation of GEFs, GAPs, and actin-binding proteins aids pathogenic mycobacteria to establish its niche within host macrophages through the disruption of actin stress fibres, actin depolymerisation, and inhibition of phagosome maturation.

### PknG manipulates host programmed cell death and inflammatory pathways to facilitate mycobacterial survival

*Mtb* infection triggers intracellular signalling pathways, enhancing pro-inflammatory responses that are crucial for controlling*Mtb* replication and the immunopathologic response (80). Apoptosis of infected macrophages is associated with diminished pathogen viability (81). However, virulent mycobacteria are known to evade apoptosis which aids the pathogenesis of these strains (82, 83).

Apoptotic chromatin condensation inducer in the nucleus (Acin1, also known as acinus) is a multifunctional protein with proposed roles in apoptosis and alternative RNA splicing (84). We identified Acin1 with differentially up-regulated p-sites S937, S656, and S425. Acin1 undergoes several proteolytic cleavages during apoptosis. However, Akt-mediated phosphorylation of Acin1 on S422 and S573 residues promotes resistance against proteolytic/apoptotic cleavage in the nucleus and inhibition of acin1-dependent chromatin condensation, thereby facilitating cell survival (85). Interestingly, Banfalvi (2014) identified anomalies in chromatin condensation associated with the apoptosis process in *Mtb*-infected macrophages (86).

Acin1 is also an auxiliary component of the exon junction complex (EJC) which is assembled throughout pre-mRNA splicing. The recruitment of a trimeric complex composed of Acin1, SAP18, and RNPS1 to the EJC was reported to modulate apoptotic and splicing regulation. Nevertheless, Acin1 independently regulates splicing profiles of apoptotic genes (87, 88). Our findings, therefore, suggest that PknG plays a role in the inhibition of apoptosis to promote bacterial survival and persistence.

Autophagy is known to contribute to the killing of intracellular microbes, including *Mtb*, by modulating host resistance against infections and controlling cellular survival (89–91). Also, autophagy activation aids in regulating inflammation, contributing to a more efficient innate immune response against *Mtb. In vitro* studies have shown that mycobacteria escaping from phagosomes into the cytosol are ubiquitinated and targeted by selective autophagy receptors (92, 93), such as Sequestosome 1 (SQSTM1; also known as p62).

We identified SQSTM1 to be differentially phosphorylated on residues S269 and S27. This signalling adaptor is central to cell survival and proliferation through the activation of the mechanistic target of rapamycin complex 1 (mTORC1). The mechanistic target of rapamycin (mTOR) is a key regulator of autophagy, cell metabolism, growth, proliferation, translation initiation, and cytoskeletal organization. Importantly, autophagy is inhibited by mTORC1 activation (94). Phosphorylation of T269 and S272 (T272 in *Mus musculus* on SQSTM1 seems to be necessary for autophagic inhibition under nutrient-rich conditions (95). Hence, SQSTM1 phosphorylation in the presence of PknG plays a role in facilitating mycobacterial survival and proliferation within the host by diminishing autophagy-mediated clearance. Our findings suggest that PknG enhances mycobacterial virulence by phosphorylating and inhibiting components of the intrinsic cell death machinery.

Moreover, during impaired autophagy, SQSTM1 accumulates and activates inflammation via nuclear factor kappa B (NF-κB) (96). We identified NF-κB1 subunit p105 to be differentially phosphorylated at S447 in the presence of PknG. NF-κB1 p105 functions both as a precursor of NF-κB1 p50 and as a cytoplasmic inhibitor of NF-κB. The consequence(s) of S447 phosphorylation has yet to be explored. However, well-studied p150 p-sites, such as S893, S903, S907, S927, and S932, point to phosphorylation being a key factor in the outcome of proteasomal processing (partial or complete degradation) of p105 (97–99). Importantly, p105 is also a negative regulator of MAPK activation downstream of various receptors, including TLRs and tumour necrosis factor (TNF) receptors (100, 101).

The mycobacterial cell wall contains several pro-inflammatory TLR2 ligands and induces activation of the MAPKs and NF-κB pathways (102, 103). The MAPK pathways play an important role in enhancing antimycobacterial activity and the production of pro-inflammatory mediators, including TNF-α (102), which contribute to phagocytosis, intracellular killing, T cell activation, and granuloma formation (104). Wu et al. (2018) demonstrated that overexpression of PknG in macrophages decreased intracellular cytokine levels, thus promoting mycobacterial survival (105). In particular, PknG was shown to inhibit the inflammatory response through suppression of NF-κB and ERK1/2 pathways. Hence, our findings of differential phosphorylation of NF-κB1 p105 potentially provide a mechanistic basis for the inhibitory role of PknG in pro-inflammatory cytokine induction.

### PknG promotes mycobacterial survival through translational control

Translational regulation of host mRNAs provides direct and rapid alterations in intracellular levels of the encoded proteins, while simultaneously providing reversibility through modifications such as phosphorylation (106). Therefore, cellular adaptation during physiological stress conditions, such as pathogen infection, attenuate general cellular translation by manipulating the activity of the protein synthesis machinery to antagonise the host immune system. Translational control occurs predominately at the initiation phase which is mainly modulated by the activity of several eukaryotic initiation factors (eIFs). We identified a total of 6 eIFs as differentially up-regulated phosphoproteins in our data, namely eIF2B5 (also known as eIF2Bε), eIF3B, eIF4G3, and eIF5B. Since eIF3B and eIF5B are both involved in IRES-dependent translation initiation (107–109), their differential phosphorylation in the presence of PknG supports the existence of non-canonical translation mechanisms that could contribute to the environmental adaptation by pathogenic mycobacteria. For example, under conditions of cellular stress and subsequent eIF2α phosphorylation (inactivation), IRES-dependent translation of X chromosome-linked inhibitor of apoptosis protein (XIAP) mRNA relies on eIF5B (107). Notably, the XIAP is associated with cell survival by protecting cells from caspase-mediated apoptosis (22–25). Additionally, the eIF4G3 protein is a central player in the initiation of translation since it also contains binding domains for mRNA, eIF4A, and eIF3. Many viruses have been shown to interfere with the function of the eIF4F complex to gain control of translation. Some viruses cleave eIF4G3 to arrest host protein translation (110) while other viruses co-opt eIF4G3 to generate viral proteins (111). Overall, the differential phosphorylation of several eIFs in our data suggests that PknG confers substantial control on protein translation and turnover in host cells, potentially preventing the activation of antimycobacterial responses.

### PknG contributes to the regulation of alternative mRNA splicing

Alternative splicing of transcripts brings unprecedented diversity to the eukaryotic proteome that can alter protein stoichiometry and physiological consequences (112). Recent studies suggest that the co-option and hijacking of host splicing machinery by pathogens contributes to the perturbation of the host immune response during infection (113, 114). Findings by Kalam et al. (2017) strongly indicated that infection with*Mtb* alters the global patterns of alternate splicing within macrophages (114). Several host splicing factors having been previously identified as interacting partners for various mycobacterial proteins (115, 116). Comparably, we identified a subset of interacting host spliceosomal proteins to be differentially phosphorylated in the presence of PknG, including U4/U6 small nuclear ribonucleoprotein Prp3 (PRP3), heterogeneous nuclear ribonucleoprotein A1 (hnRNP-A1), THO complex subunit 4 (THOC4), and U1 small nuclear ribonucleoprotein 70 kDa (snRNP70). Differential levels of these phosphoproteins in our data suggest that PknG promotes aberrant interactions within the host splicing machinery to interfere with the ability of the macrophage to kill the mycobacterium.

Moreover, the co-transcriptional RNA processing enzyme, 5′-3′ exoribonuclease 2 (XRN2) was differentially phosphorylated on residues S499 and S501 in the presence of PknG. XRN2 plays a major role in ribosomal RNA (rRNA) maturation, gene silencing (117–120), and DNA damage response and replication stress (121). Cells deficient in XRN2 displayed increased double-strand breaks (DSBs) and genomic instability, along with decreased DNA repair capacity (121). Interestingly, Castro-Garza et al. (2018) demonstrated that *Mtb* infection promoted genomic instability in macrophages through the observation of several DNA damage markers (122). Another study showed that *Mtb* induced DSBs, compared to single-stranded DNA generation initiated by the avirulent strain (123), which provided a survival niche through activation of the ATM-Akt signalling cascade that results in the inhibition of apoptosis and accentuation of cell growth.

## Conclusion

In this study, we have employed label-free mass-spectrometry based phosphoproteomics to unravel the mechanisms that pathogenic mycobacteria use – particularly, those mediated by PknG – to evade host immune responses and survive during the early stages of infection. Our data revealed the multifactorial signalling pathways exploited by PknG to promote mycobacterial survival in host macrophages, including perturbation of host actin cytoskeleton organisation and phagosome maturation, inhibition of apoptotic pathways and NF-kB-mediated inflammatory responses, impairment of autophagy, as well as manipulation of alternative splicing, translation and of DNA repair processes. In each case, we find supporting data in the literature validating these perturbations, but importantly our phosphoproteomic data provides significant new mechanistic detail on PknG-mediated post-translational control of these host processes.

This work, therefore, considerably increases our current knowledge of mycobacterial pathogenicity and may additionally identify novel molecular approaches to effectively reactivate particular host cellular pathways to restore macrophage capability of mycobacterial elimination. However, to confidently classify these proteins as direct or indirect host substrates of PknG, validation studies are necessary. Validation through targeted parallel reaction monitoring (PRMs) MS assays and imaging experiments are now underway and will be reported elsewhere.

## Acknowledgements

We would like to thank Professor Jean Pieters for graciously donating the bacterial strains used in this study. This work is based on the research supported, in part, by the National Research Foundation (NRF) of South Africa (Grant Numbers: 467126 and 95984). S.S.B. thanks the NRF for a Master’s bursary. K.C.N. thanks the NRF and UCT/CSIR for PhD bursaries. D.L.T. was supported by the South African Tuberculosis Bioinformatics Initiative (SATBBI), a Strategic Health Innovation Partnership grant from the South African Medical Research Council and South African Department of Science and Technology. N.C.S. thanks the South African Medical Research Council for a Junior Research Fellowship. J.M.B. thanks the NRF for a South African Research Chair grant.

